# Purification of the mammalian NgBR/hCIT *cis*-prenyltransferase complex: Identification of a conserved carboxyterminal RxG motif crucial for enzymatic activity

**DOI:** 10.1101/139675

**Authors:** Kariona A. Grabińska, Ban H. Edani, Eon Joo Park, Jan R. Kraehling, William C. Sessa

## Abstract

*Cis*-Prenyltransferases (*cis*PTs) constitute a large family of enzymes conserved during evolution and present in all domains of life. In eukaryotes and archaea, *cis*PT is the first enzyme committed to the synthesis of dolichyl-phosphate (DolP). DolP is obligate lipid carrier in protein glycosylation reactions in mammals. The homodimeric bacterial enzyme, undecaprenyl diphosphate synthase (UPPS) generates 11 isoprene units and has been structurally and mechanistically characterized in great detail. Recently our group discovered that unlike UPPS, mammalian *cis*PT is a heteromer consisting of NgBR (NUS1) and hCIT (DHDDS) subunits and this composition has been confirmed in plants and fungal *cis*PTs. Here, we establish the first purification system for heteromeric *cis*PT and show that both NgBR and hCIT subunits function in catalysis and substrate binding. Finally, we identified a critical RxG sequence in the C-terminal tail of NgBR that is conserved and essential for enzyme activity across phyla.

## INTRODUCTION

Dolichyl phosphate is an obligate lipid carrier for protein N-glycosylation, 0-mannosylation, C-mannosylation and GPI-anchor synthesis in eukaryotic cells and undecaprenyl phosphate is essential for peptidoglycan biosynthesis in bacteria. *cis*-Prenyltransferase (*cis*PT) is the rate limiting enzyme committed to dolichyl phosphate biosynthesis in Eukaryotes, and Archaea as well as undecaprenyl phosphate biosynthesis in Eubacteria (1,2). Both eukaryotic and prokaryotic *cis*PTs belong to a large protein family, well conserved during evolution (1) and the *cis*PT family was identified amongst the 355 protein families that trace to the last universal common ancestor of all cells (LUCA, or the progenote) by phylogenetic criteria (3).

The bacterial enzyme, undecaprenyl diphosphate synthase (UPPS) is a homodimeric enzyme that catalyzes chain elongation of farnesyl diphosphate (FPP) by sequential reactions with eight isopentenyl diphosphate molecules (IPP). UPPS has been structurally and mechanistically characterized in great detail (4,5) however, the eukaryotic enzyme has not been purified to date. Our group discovered that unlike UPPS, mammalian and fungal *cis*PT is heteromeric complex consisting of NgBR (Nus1) and hCIT (DHDDS) subunits in human cells and these findings were confirmed for a number of plant *cis*PTs (6–10) demonstrating a major difference in the composition of *cis*PT activity in prokaryotes and eukaryotes. Moreover, loss of function mutations identified in patients via exome sequencing in either NgBR or hCIT causes a congenital glycosylation disorder, and results in severe clinical manifestations including cognitive defects and retinitis pigmentosa (11–15) and microdeletions within NgBR locus are linked to pediatric epilepsy(16,17). Further phylogenetic analysis of NgBR and UPPS suggests that a C-terminal motif important for mammalian and fungal *cis*PT is shared by the single subunit *cis*PT UPPS (1).

In the present study, we have purified and characterized for the first time a heteromeric *cis*PT complex composed of NgBR and hCIT subunits. Biochemical characterization of purified wild-type (WT) and various mutants of the NgBR/hCIT complex shows that both subunits contribute to catalytic activity. Furthermore, we provide evidence that a conserved C-terminal motif -RxG-motif is critical for enzyme catalysis in both two-component and single-subunit *cis*PTs.

## RESULTS

### Purification and biochemical characterization of human *cis*-PT

Although NgBR and hCIT or its orthologs are essential subunits for *cis*PT activity in mammals, fungi and plants, the heteromeric complex and its activity have not been isolated and purified. Thus, we developed a purification scheme to isolate the NgBR/hCIT complex (h*cis*PT). Optimal expression and tagging strategy was first established using a previously described expression system in a triple knockout strain lacking *nus1Δ rer2Δ srtl Δin S*. *cerevisiae* (13). Cell survival was ensured due to expression of Gl*cis*PT on the *URA3* plasmid and cells were co-transformed with the *LEU2* and *MET15* plasmids bearing WT or epitope tagged versions of hCIT and NgBR respectively. The yeast cells were streaked onto complete plates or synthetic complete medium containing 1% 5-fluoroorotic acid (FOA). Since Ura3 protein, which is expressed from the *URA3* marker present in the plasmids, converts FOA to toxic 5-fluorouracil, the survival of the yeast cells on the FOA plate depends on the functionality of the expressed NgBR/hCIT complex. hCIT isoform 1 (UniProtKB# Q86SQ9-1) and isoform 2, (UniProtKB # Q86SQ9-2), but not isoform 3 (UniProtKB# Q86SQ9-3) supported the growth of yeast when co-expressed with NgBR in the triple delete strain (data not shown). N- terminal tagging, but not C-terminal tagging of hCIT, generated a construct indistinguishable from wild-type (WT), non-tagged hCIT in yeast. Both N- and C- terminal tagging of NgBR reduced its stability and activity when expressed in yeast, therefore a 6-His tag was placed internally after Gly31, between the putative signal anchor and TM1 (Figure 1A). This tag did not affect the growth phenotype and *cis*PT activity in yeast co-expressing N-terminally Strep-tagged hCIT (not shown).

**Figure 1.**
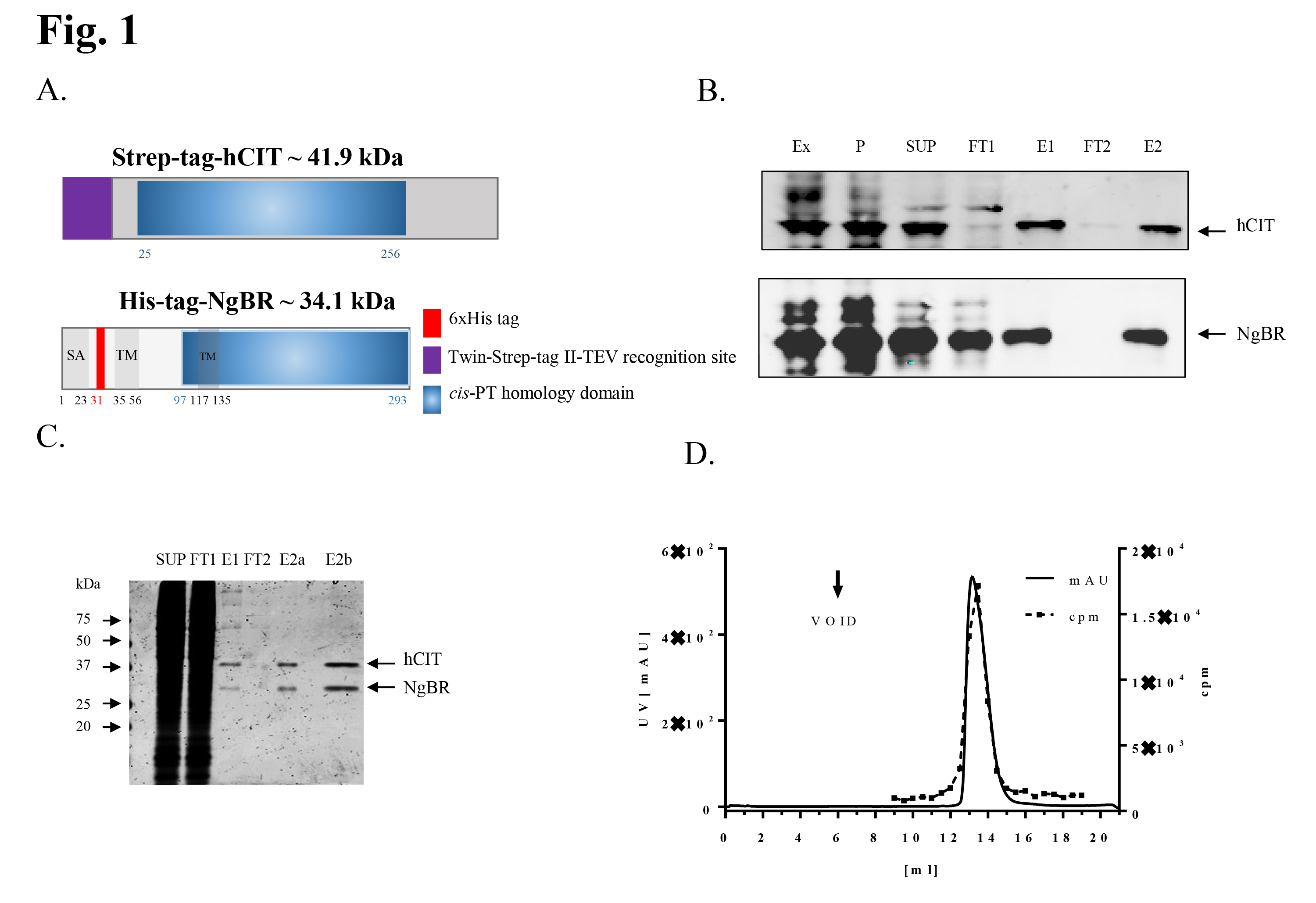
Purification of NgBR/hCIT *cis*-PT complex. **A**. Epitope tagged NgBR and hCIT were transiently expressed in Expi293 cells from bicistronic vector and purified as decribed in the text (SA, putative signal anchor; TM, putative transmembrane domain). **B** is a representative western blot monitoring purification of NgBR/hCIT complex (Ex-Total Crude Extract, P-200,000g membrane pellet, SUP- 200,000g supernatant after Triton X100 solubilization of the membrane fraction, FT1 and FT2- flow through from Strep-Tactin and Nickle column respectively, E1-elution from the Strep-tactin and E2- final elution from the Nickle column). **C** is a Coomassie stained SDS/PAGE showing SUP, FT1, E1, two (E2a) and eight times concentrated final fraction(E2b). **D** shows size exclusion chromatography profile of purified NgBR/hCIT *cis*-PT complex. UV absorption at 254 urn was measured as a readout for protein elution (lefty axis, black line). Incorporation of C^14^ labeled IPP into organic fraction was measured as a readout for *cis*-PT activity (right y axis, red line).

Next, the cDNAs encoding 6xHis-NgBR and Strep-tagged-hCIT (Fig1A) were cloned into an IRES-containing bicistronic vector allowing the simultaneous expression of two proteins separately but from the same RNA transcript. This was necessary based on our previous work demonstrating that co-translation of both subunits was required for activity in an in vitro translation system (13). To generate protein for purification, the construct was transiently transfected into Expi293F cells and cells collected 72 hours later by centrifugation. The pellet was washed with PBS, and lysed in detergent free buffer (Ex). To pre-fractionate cell extracts prior to affinity chromatography, crude membrane extracts were subjected to ultracentrifugation, and pellet (P) solubilized by Dounce homogenization in the presence with 0.5% Triton X100, and solubilized protein cleared by a second ultracentrifugation step (SUP). N-terminally Strep-tagged hCIT and internally tagged 6xHis-NgBR complex was purified using a dual affinity purification scheme **(Figure 1B; Table I).** Each step of purification was monitored by Western Blotting and measurement of *cis*PT activity and this scheme enriched the specific activity of the enzyme 1,700 fold over the starting material. As judged by SDS-PAGE and Coomassie staining of each fraction, the hCIT/NgBR complex was approximately 95% pure **(Figure 1C)** and the purified heteromeric complex ran as monodispersed peak on size exclusion chromatography **(Figure 1D; black line)** that tracked with *cis*PT activity **(Figure 1D; dashed line)**.

The purified hCIT/NgBR complex required Mg^2+^ ions for its activity consistent with other studies of the bacterial and eukaryotic enzymes (18–24) with maximum activity at 0.5-2mM MgCl_2_. **(Figure. 2A)**. The purified enzyme had a broad pH optimum **(Figure 2B)** (19) and was highly stable demonstrating linear *cis*PT activity occurred over 24hrs at 37°C **(Figure 2C)**. To test the influence of lipids on human *cis*PT activity, Triton X100 was removed from the enzyme preparation by buffer exchange and activity was measured upon the addition of Triton X100 or different phospholipids at concentration above their critical micellar concentration. As seen in **Figure 2D**, phosphatidic acid modestly increased enzyme activity (1.8 fold) whereas other phospholipids had a marked effect in activating of the enzyme. Cardiolipin increased *cis*PT activity over 7 fold and phosphatidylcholine, phosphatidylethanolamine, phosphatidylinositol and phosphatidylserine increased activity by approximately 8–12 fold suggesting that the lipid environment strongly impacts *cis*PT activity.

**Figure 2.**
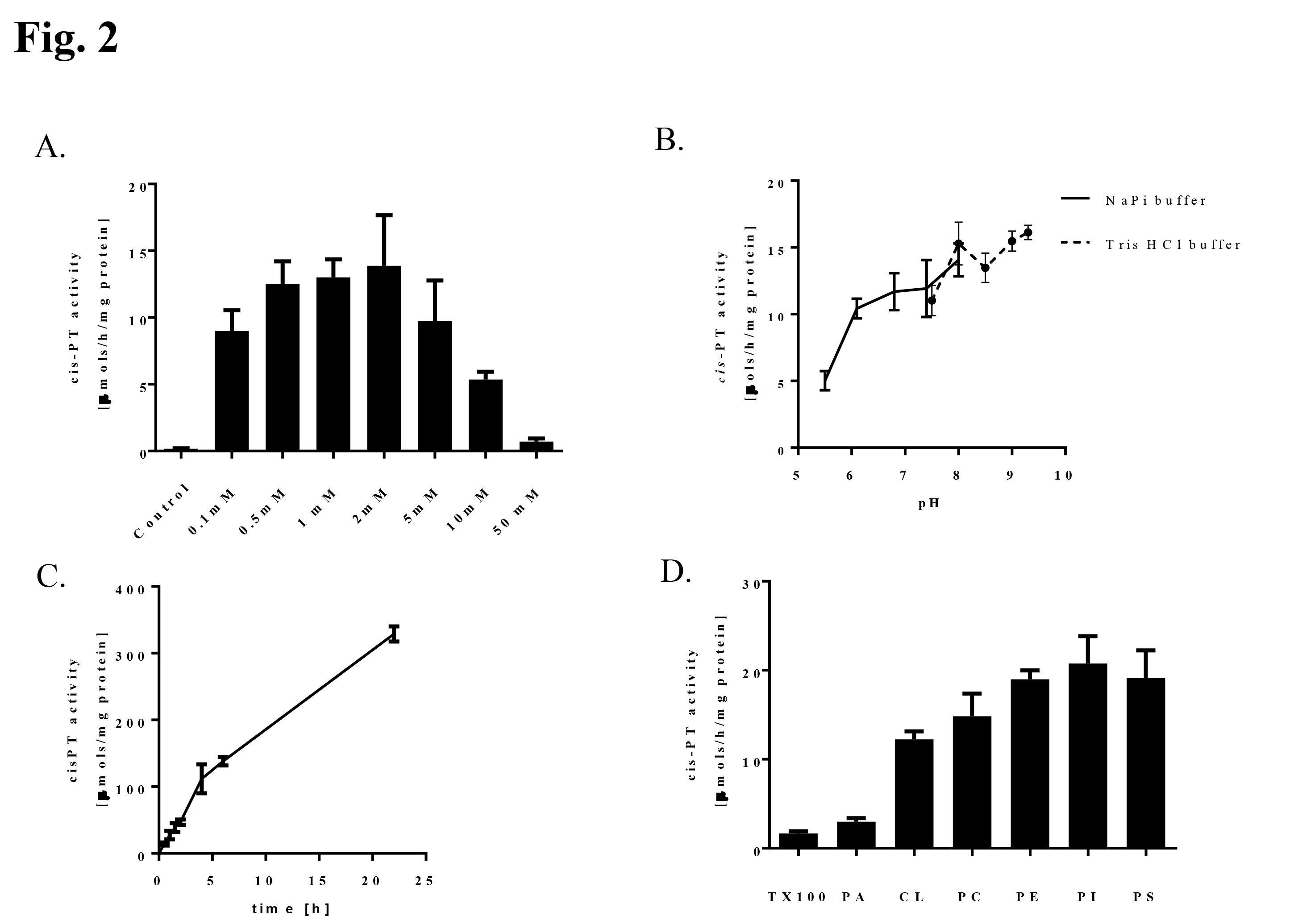
Biochemical properties of purified human *cis*-PT. *cis*-PT activity was measured using purified h*cis*-PT complex as described in Material and Methods unless otherwise stated. **A**. Optimal concentration of MgCl_2_. *cis*-PT activity was measured in the presence of 5mM EDTA (to chelate MgCl_2_ in the enzyme storage buffer,) or 0.1, 0.5, 1, 2, 5, 10 and 50 mM MgCl_2_. B. pH dependence of *cis-PT* activity. *cis-PT* activity was measure in the 50 mM sodium phosphate buffer (pH 5.5; 6.1;7.4; 8.0) or 50 mM Tris HCl (pH 7.5; 8.0; 8.5; 9.0; 9.3). C. Time dependence of *cis-PT* activity. D. Phospholipids stimulate *cis-PT* activity. *cis-PT* activity was measured in the buffer containing 0.35% (w/v) Triton X100, 1% (w/v) Phosphatidic acid (PA), 1% (w/v) Cardiolipin (CL), 1% (w/v) Phosphatidylcholine (PC), 1% (w/v) Phosphatidylethanolamine (PE), 1% (w/v) Phosphatidylinositol (PI) or 1% (w/v) Phosphatidylserine. Data are mean ±S.D values of three technical replicates,

### Protein Sequence Alignment of *cis*-prenyltransferase homology domain bearing proteins

Sequential alignment of nonredundant homomeric *cis*PTs from all three domains of life together with heteromeric orthologs define the position of *cis*PT homology domain among NgBR orthologs **(Figure 3)**. Analysis of the architecture of representative *cis*PT homology domains revealed number of characteristic features for each of the group (single-subunit orthologs, hCIT orthologs and NgBR orthologs) separately or shared between two groups (1,4,8,25,26). Only four of the previously identified five conserved regions (4) are present in all three groups. The NgBR group lacks Region II and has highly degenerate Regions I and III. Among other conserved residues, the NgBR class is missing a highly-conserved stretch of the six amino acid residues involved in FPP binding and catalysis in Region I (boxed region) found in UPPS and hCIT. Regions IV and V, including the predicted dimer interface based on the structural data obtain for the UPPS of *M luteus* and *E. coli* (27,28) are well conserved amongst all three groups. Finally, the NgBR group shares with homomeric enzymes, but not with the hCIT group, an -RxG C-terminal conserved motif (see red arrowheads)(1,13) with G being absolutely conserved in EcUPPS and NgBR class. The R residue is substituted with N in SaUPPS and a majority of plant and fungal NgBR orthologs (6–10,13,29). In a majority of the cases, X is any nonpolar amino acid however putative *Hypocreales* NgBR orthologs as well as UPPS of some *Trypanosomatidae* have positively charged H or R at this position. Meta-analysis of available experimental data concerning *cis*PTs, suggest that both the catalytic motif of Region I and the C-terminal -RxG-motif are indispensable for enzymatic activity being part of the same subunit or separated between hCIT (catalytic motif) and NgBR (RxG) class of *cisPTs* proteins.

**Figure 3.**
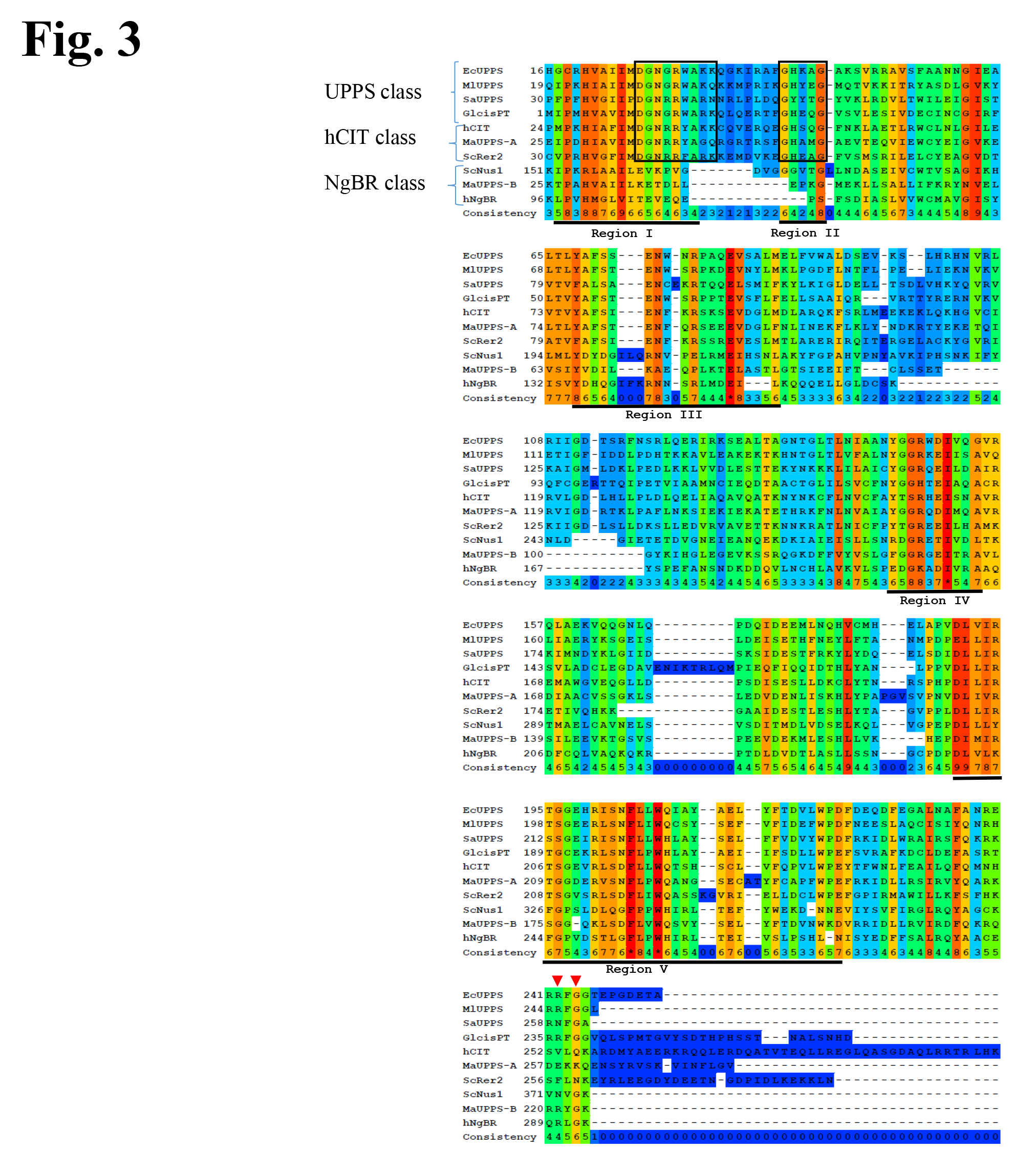
Multiple aligmuent of the proteins bearing *cis*-PT homology domain. Proteins represented in the figure are: single subunit *cis*-PTs: EcUPPS (*Escherichia coli*, GenBank P60472), M1UPPS (*Micrococcus luteus;* GeneBank BAA31993.1), SaUPPS (*Sulfolobus acidocaldarius;* GenBank WP_011277635.1) and GlcisPT (*Giardia lamblia;* GenBank XP_001709868.1); orthologes of hCIT *cis*-PT subunit: hCIT (human, GenBank BAB14439), ScRer2 (*Saccaraomyces cerevisiae*, GenBank P35196), MaUPPS-A (*Methanosarcina acetovorans*, GenBank AAM07075.1); orthologes of NgBR *cis*-PT subunit: NgBR (human, GenBank NP_612468), ScNus1 *(Saccaraomyces cerevisiae*, GenBank NP_010088), MaUPPS-B *(Methanosarcina acetovorans*, GenBank) WP_011024281.1). The conservation scoring is performed by PRALINE. The scoring scheme works from 0 for the least conserved alignment position, up to 10 (*)for the most conserved alignment position. Position of five conserved regions originally described among *cis*-PTs is underlined. Residues involved in catalysis and substrate binding conserved between hCIT orthologues and single subunit *cis*-PTs are boxed. Red triangles indicate position of conserved Arginine/Asparagine and Glycine in C-terminal -RxG- motif of NgBR orthologues and homomeric *cis*-PTs. Black triangle indicates position of strictly conserved C-terminal Phenylalanine/Tyrosine in *cis*-PT homology domain.

### Phenotypic analysis of *hcis*PT mutants in yeast and biochemical characterization of purified proteins supports importance of RxG motif and position of *cis*PT homology domain in NgBR

To examine the role of the conserved RxG motif in h*cis*PT, and to test the predicted position of the *cis*PT homology domain in NgBR, we performed phenotypic analysis of a series of truncation and point mutants of NgBR or hCIT using the above mentioned yeast triple deletion strain *rer2Δ srt1Δ* and *nuslΔ (3)*. No growth defects were observed with complexes containing NgBR^H100A^ (His the first amino acid in Region I and is highly conserved among proteins bearing *cis*PT homology domain), NgBR^R290H^ (a human mutation causing a glycosylation disorder), Δ85-NgBR (truncated protein lacking first 84 amino acid including TM domain conserved across eukaryotic orthologs of NgBR) or hCIT^K42E^ (a human mutation causing retinitis pigmentosa) in comparison to WT *cis*PT composed of NgBR/hCIT (Figure 4A). Essentiality of the conserved D^34^ in hCIT corresponding the D^26^ (18,27,30) of EcUPPS was confirmed since the hCITD^34A^/NgBR complex did not support the growth of triple knockout yeast cells. Cells expressing truncated Δ101-NgBR (lacking the first 100 amino acids) had a severe growth defect suggesting that the first conserved region in NgBR, despite its degeneration, was important for function. This finding is in line with the recent structural analysis of plant *Z,Z-* famesyl diphosphate synthase (zFPPS) suggesting that N-terminus might be able to extend to the active site of the neighboring monomer near the C-terminus as an important functional regulator (31).

**Figure 4.**
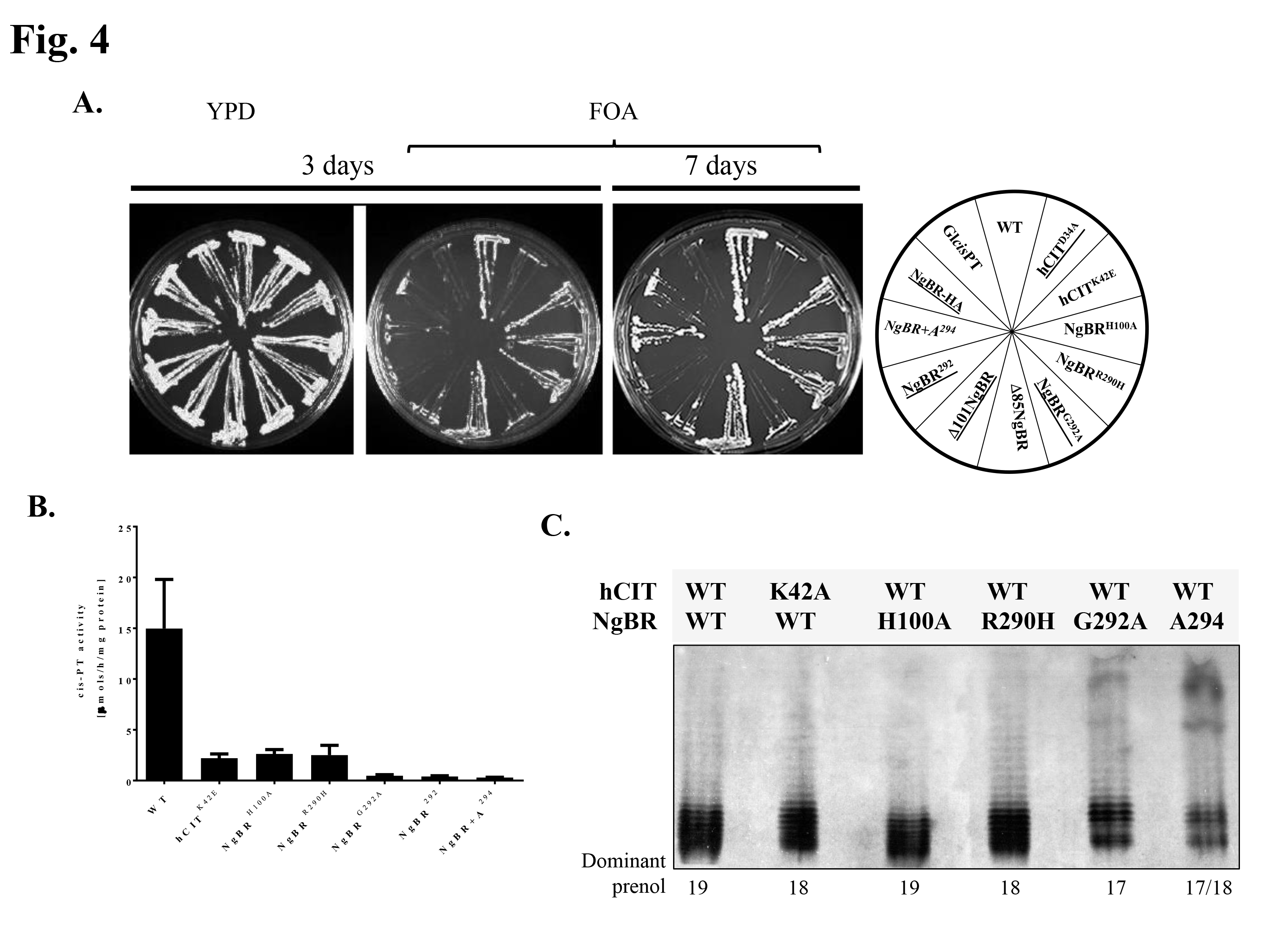
Characterization of human *cis*-PT mutants. **A**. The *rer2Δ, srt1Δ, nus1Δ* triple deletion strain expressing *G. Iamblia cis*-PT from *URA3* plasmid was co-transformed with the *LEU2* and *MET15* plasmids bearing wild or mutated variant of hCIT and NgBR as indicated. The cells were streaked onto complete plates (YPD) or synthetic complete medium containing 1% 5-fluoroorotic acid (FOA). The Ura3 protein, which is expressed from the *URA3* marker present in the plasmids, converts FOA to toxic 5-fluorouracil. The growth of cells was monitored overtime to assess phenotypic differences. The combination of alleles affecting the growth is indicated in italic and the combination not supporting the growth is marked in red. **B.** *cis*-PT activity was measured using purified WT or mutant h*cis*PT complex as described in Material and Methods. Mean ±S.D values of 3-9 independent measurements from at least 2 independent isolations of enzyme except for hCIT/NgBR+A^294^ and hCIT/NgBR^292^ performed on one batch.). **C**. Reverse-phase TLC of dephosphorylated polyprenols formed by h*cis*PT mutants. Equal amounts of product were analyzed to reveal quality differences.

Finally, conservation of the RxG motif was confirmed by the series of mutants leading to severe growth defects. Deletion of the C-terminal K^293^ (NgBR^292^), the addition of A to NgBR (NgBR+A^294^), HA tagging (NgBR-HA) or *NgBRaznA* substitutions all reduced growth documenting the importance of the last 4 residues in NgBR for function **(Figure 4A)**. Next, we compared the steady state activities of purified mutants able to support the growth of yeast to the purified WT enzyme. We did not include the N-terminal truncation variants of NgBR since introducing the epitope tag after the predicted TM1 (35-56aa) rendered the enzyme inactive in yeast. As shown in **Figure 4B**, all mutants analyzed displayed lower enzymatic activity compared to WT enzyme and mutations that strongly impacted growth in yeast (NgBR^G292A^, NgBR^292^ and NgBR+A^294^) had the lowest enzymatic activity varying co-substrates FPP and IPP **(Figure 4- supplemental figure 1)**. Next, the polyprenol reaction products were examined by TLC. Since the mutants had reduced activity, no products were detected on the TLC plate when equal amounts of reaction products were loaded from WT enzyme compared to the mutants, therefore an overload of products was used to examine the chain length of prenols generated from the different enzymes. As seen in **Figure 4C**, WT NgBR and NgBR^HlOOA^ generated dominant prenols of C_95_ (19 units), disease mutants, hCIT^K42E^ and NgBR^R290H^ complexes generated prenols of C_90_ (18 units) while NgBR^G292A^ and NgBR+A^294^ yielded polyprenols that were one or two units shorter. This data support the important role of C-terminus, in particular G^292^ in *cis*PT activity.

To further characterize h*cis*PT and elucidate the impact of mutations on catalysis and substrate binding, steady state kinetic parameters were measured for complexes containing the WT subunits, hCIT/NgBR, hCIT^K42E/^NgBR, hCIT/NgBR^H100A^, hCIT/NgBR^R290H^ and hCIT/NgBR^G292A^ (Table 2). The K42E substitution of hCIT reduced k_cat_, and increased the K_m_ for FPP but not the affinity for IPP. NgBR^H100A^ and NgBR^R290H^ decreased the kcat around 4-fold compared to WT enzyme. However, NgBR^H100A^ did not exhibit marked changes in affinity for IPP or FPP, whereas NgBR^R290H^ had moderately but significantly reduced affinities for IPP. Substitution of G292A in NgBR, which markedly reduced the growth phenotype in yeast, decreased the catalytic activity of the enzyme to an even greater extent with kcat that is 10-fold lower than the WT. In this case, there were modest (5-fold) but significant differences in the K_m_ for IPP, and a slight decrease in the K_m_ for FPP.

### The RxG motif in homomeric EcUPPS and GlcisPT is critical for *cis*PT function and enzymatic activity

Comparison of the primary amino acid sequences of single- and two-component enzymes reveals that both classes share a conserved RxG C-terminal motif (see Figure 3). Based on the crystal structure of UPPS, R^242^ in the *RXG* motif is involved in the binding of the diphosphate group of iPP (30). Further, the role of C terminus in IPP binding and catalysis is supported by structural information obtained for decaprenyl diphosphate synthase of *M tuberculosis* and recent studies on zFPPS of *Solanum habrochaites* (31,32) implicating the C terminus in IPP binding. To verify importance of RxG motif in homomeric *cis*PTs experimentally, we assayed growth in the triple deletion strain in yeast, transformed with WT or mutant forms of homomeric forms of *cis*PT, namely EcUPPS and Gl*cis*PT. Cells were transformed with *LEU2* plasmids bearing: *WT, R242H* and *G244A* of EcUPPS or *WT*, *R236H*, and *G238A* of Gl*cis*PT and phenotypes scored over 7 days. As seen in Figure 5A, WT *E.coli* and *G. lamblia cis*PTs exhibited normal growth phenotypes on YPD and FOA plates and R to H mutants for each construct had similar phenotypes compared to WT transformed cells. In contrast, UPPS^G244A^ and GlcisPT^G238^Amutants failed to grow on FOA plates demonstrating an essential role of G in the RxG motif in homomeric *cis*PT function. To biochemically support the phenotypic data, *cis*PT activity was measured using purified WT, R236H and G238A mutants of GlcisPT. Mutation of both residues reduced activity, with *Glcis*PT^236^ having approximately 5 times lower activity and Gl*cis*PT^G238A^ being virtually inactive. To verify whether the G238A mutant is enzymatically inactive or generates shorter polyprenols an overload of products was used to examine the chain lengths of undecaprenol from the different enzymes by TLC. As seen in Figure 5C, WT enzyme generated undecaprenyl (C_55_ or 11 IPP units), Gl*cis*PT^R236H^ yielded polyprenols that were one unit shorter compared to WT enzyme (C_55_ or 11 IPP units) and no product was observed for Gl*cisPT^G238A^* in agreement with its undetectable enzymatic activity and the phenotypic data.

**Figure 5.**
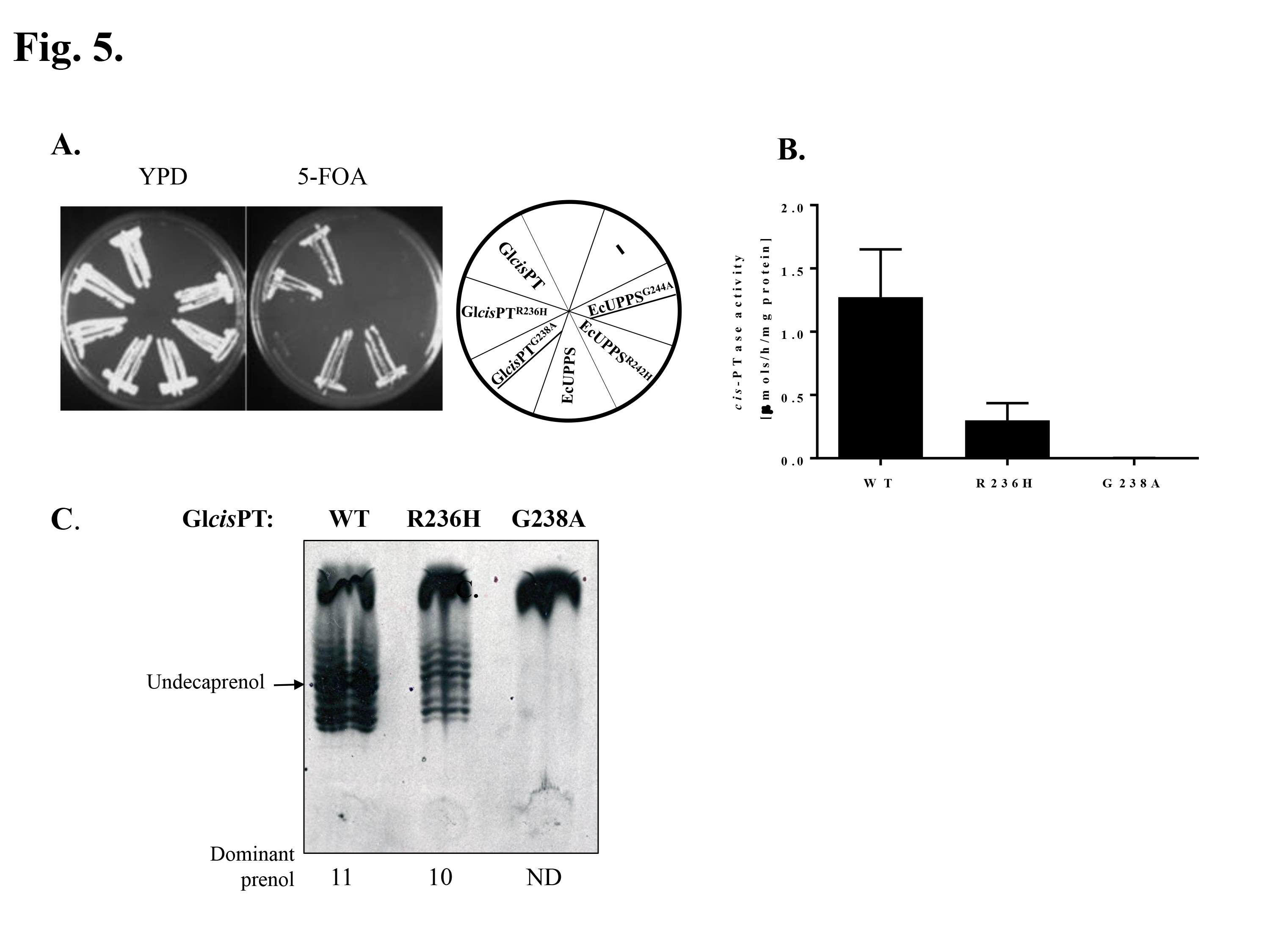
Characterization of EcUPPS and Gl*cis*-PT mutants in RxG motif reveals is essentiality. **Panel A**. The rer2Δ, srt1Δ, nus1Δtriple deletion strain expressing G. lamblia *cis*-PT from *URA3* plasmid was transformed with the *LEU2* plasmid bearing WT or mutants ORFs for Gl*cis*PT or EcUPPS. As a negative control, cells were transformed with empty vector. The mutated genes G244A and G238A did not supporting the growth and are underlined. **B**. *cis*-PT activity was measured using purified wild type or mutant Gl*cis*-PT. Data are mean ±S.D values of three technical replicates. **C**. Reverse-phase TLC of dephosphorylated polyprenols formed by purified Gl*cis*-PT.

### Presence of heteromeric *cisPT* in *Methanosarcina acetovorans* supports the importance of RxG motif

Heteromeric *cis*PTs was predicted in subgroup of *Euryarchaeota*, based on the fact that *Halomebacteria* and *Archaeoglobaceae* have at least two UPPS orthologs: a putative *cis*PT closely related to single subunit *Archea* enzymes, but lacking C-terminal RxG motif, and a NgBR/Nus1like protein (1,33) (Fig. 3). To test the conservation of this motif, we cloned putative undecaprenyl diphosphate synthase of *M. acetovorans* consisting of MaUPPS-A (MA3723, hCIT group) and MaUPPS-B (MA4402, NgBR/Nus1 group). The triple deletion strain bearing GlcisPT on the *URA3* plasmid was co-transformed with the *LEU2* and *MET15* plasmids expressing wild or mutated variants of MaUPPS-A and MaUPPS-B respectively. The growth of yeast cells on the FOA plates was monitored over 5 days. As seen in **Figure 6A**, co-expression of MaUPPS-A and MaUPPS-A is indispensable to support cell growth and neither, MaUPPS-A or MaUPPS-B alone are sufficient. Similarly, as it was observed in case of NgBR, mutation of His^29^ (corresponding to His^100^ of NgBR) and Arg^221^ (corresponding to Arg^290^ of NgBR) in MaUPPS-B does not impair growth, but MaUPPS-B^G223A^ substitution caused a severe growth delay. Quantification of the *cis*PT activity reveals that each of the analyzed mutation inhibits enzyme activity with most profound effect of MaUPPS-B^G223A^ substitution. In accordance with the data on both homomeric and heteromeric enzymes, mutation of the RxG motif negatively influences the chain length of the final prenol product **(Figure 6C)**.

**Figure 6.**
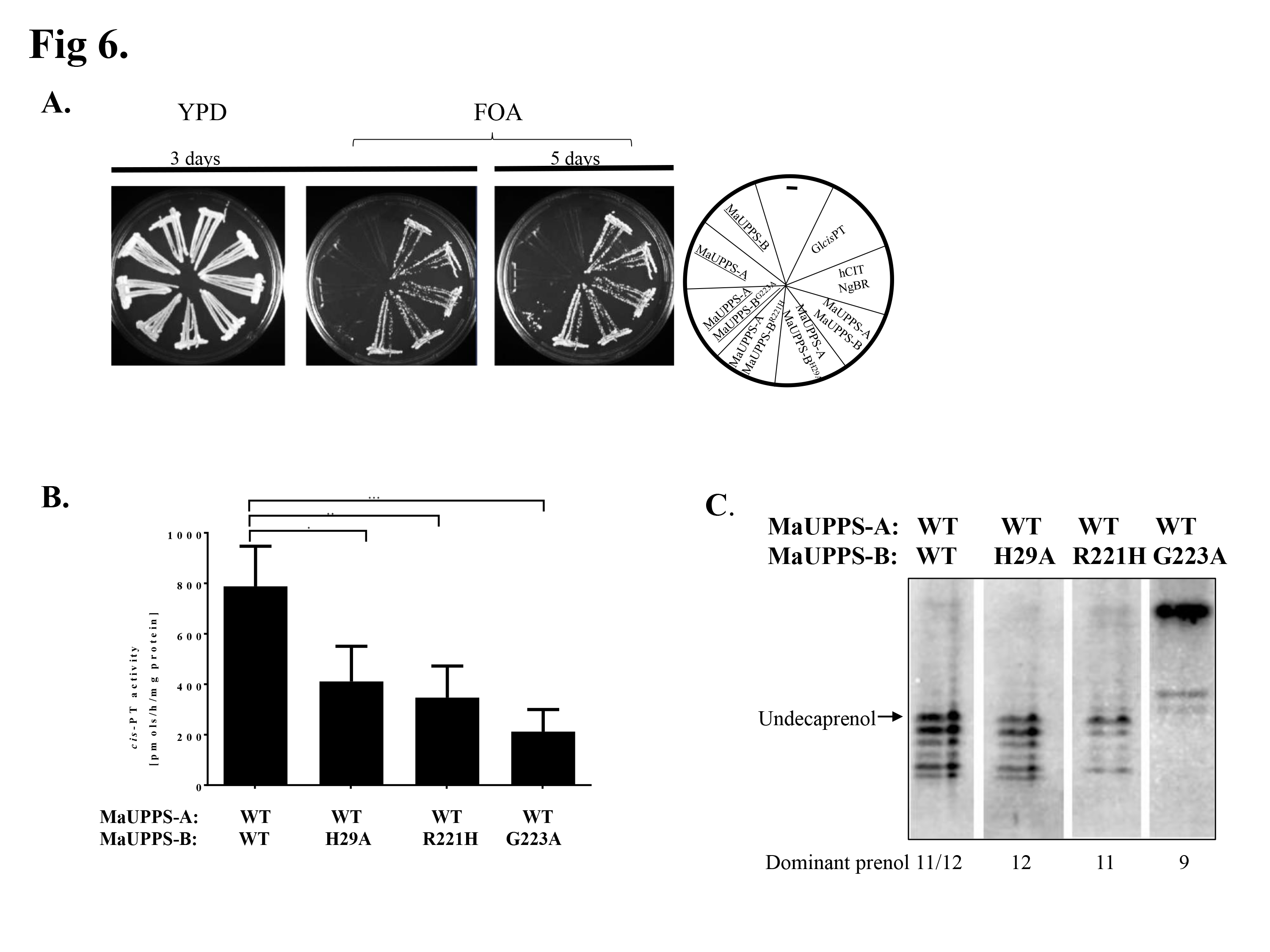
Phenotypical and biochemical analysis of heteromeric MaUPPS supports predicted length of *cis*-PT domain of NgBR and importance of C-terminal RxG motif. **A.** The *rer2Δ, srt1Δ, nus1Δ* triple deletion strain expressing *G. lamblia cis*-PT from *URA3* plasmid was co-transformed with the *LEU2* and *MET15* plasmids bearing wild or mutated variant of MaUPPS-A (hCIT/Rer2/Srt1 orthologue) and MaUPPS-B (NgBR/Nus1 orthologue) as indicated. Cells transformed with empty plasmid were used as negative control and cells transformed with *LEU2* GlcisPT or co transformed with hCIT (LEU2) and NgBR (MET15) were used as positive control. The cells were streaked onto YPD or synthetic complete medium containing 1% 5-fluoroorotic acid (FOA). The growth of cells was monitored overtime to assess phenotypic differences. The combination of alleles affecting the growth is indicated in italics and the combination not supporting the growth is underlined **B.** *cis*-PT activity was measured using *S cerevisiae rer2Δ srt1Δ nus1Δ* membrane fractions expressing WT or mutated MaUPPs subunits as indicated. Data are mean ±S.D values of 4-5 replicates in two independent experiments‥ **C.** Reverse-phase TLC of dephosphorylated polyprenols formed by heteromeric wt or mutaed MaUPPS expressed inS *cerevisiae rer2Δ srt1Δ nus1Δ* mutants.

## DISCUSSION

Here we demonstrate that two subunits of the human *cis*PT, NgBR and hCIT, are required to form a functional enzyme. The purification of the *cis*PT complex supports prior work showing that co-translation of both subunits is required for polyprenol synthesis in vitro and for survival in yeast lacking orthologs of each component(B). Moreover, the catalytic D^34^ in hCIT and RxG motif in NgBR, both conserved in homomeric *cis*PT such as UPPS, are critical for catalytic activity of the complex. These data highlight the evolutionary conservation of essential elements required for *cis*PT function throughout all walks of life.

Eukaryotic *cis*PTs were initially presumed to be homomeric based on detailed studies in undecaprenyl diphosphate synthases of *E. coli* and *M. luteus* (4,5,34–36). Recent work by us (13,25,26) and others have shown the essential role of NgBR (and its orthologs including Nus1 in *S. cerevisiae*) and hCIT (and its orthologs including Rer2 and Srt1 in *S. cerevisiae*) as both being required for *cis*PT activity and polyprenol synthesis (6,8–10,13). The expression of only hCIT or only NgBR does not support growth in the *nus1Δ, rer2Δ, srtlΔ* strain of *S*. *cerevisiae* and in vitro translation of either subunit is not catalytically active (13). At first glance, this may appear to be in conflict with previous reports showing that eukaryotic hCIT orthologs heterologously expressed in *S. cerevisiae* were active without co-expression of NgBR/Nus1 orthologs. However, this can be explained due to the presence of the endogenous Nus1 in *S*. *cerevisiae* (7,10,33–35,37–40). Furthermore, a number of hCIT orthologs in plants and protists that were studied in yeast are single-subunit enzymes that have been acquired through horizontal gene transfer, and are more closely related to EcUPPS than to hCIT (1,41). This complexity can be overcome and simplified by complementation experiments in the *nus1Δ, rer2Δ, srtlΔ* strain as performed previously (9,13) where it is clear that both NgBR and hCIT or a single-subunit enzyme is required.

Previous experiments have shown that co-expression of NgBR with hCIT results in their interaction and stabilization based on co-immunoprecipitation experiments (13,26). Whether NgBR/Nus1 serves as an accessory subunit for docking the complex to the ER membrane or is a structural component contributing to catalysis and substrate binding was not known. This idea of NgBR being a docking component was recently reported by Yamashita and co-workers (6) who identified a three-component system composed of HRBP (a NgBR ortholog), HRT (a hCIT ortholog) and rubber elongation factor (42) all of which were required for long chain prenol or natural rubber synthesis when reconstituted into washed rubber particles. Sequential alignment of nonredundant proteins bearing *cis*PT homology domains presented in this paper enabled the observation of a highly conserved C-terminal RxG motif shared between NgBR/Nus1 and the UPPS group of enzymes. Furthermore, meta-analysis of available experimental data concerning *cis*PT suggests that the C-terminal RxG motif is indispensable for enzymatic activity. The importance of the RxG motif is strengthened by the fact that NgBR^R290H^ mutation causes a congenital disorder of glycosylation due to defects in dolichol synthesis (13). To further elucidate the role of NgBR in h*cis*PT activity, we successfully purified the WT NgBR/hCIT complex and its mutated versions to homogeneity and biochemically characterized the proteins. Mutations in either subunit of h*cis*PT reduces enzymatic activity in comparison to the WT enzyme and these results are in agreement with the predicted function of the analyzed residue in homomeric *cis*PTs.

The first observation supporting a functional role of the extreme C-terminus of NgBR regulating enzyme activity is derived from patients harboring an R290H mutation in NgBR which reduces activity in vivo and in vitro (13) and now seen with the purified complex (Table 2) by 70-80%. In addition, epitope tagging the C -terminus, of NgBR reduces activity (Figure 4) and this has been previously shown in the CPT-like (CPTL) NgBR ortholog of *L. sativa* (8). Crystallographic data obtained for homomeric EcUPPS, shows that Arg^242^ in the RxG motif is involved in Mg^+2_^ IPP binding (30). In addition, the role of the C terminus in IPP binding and catalysis is supported by structural information obtained for decaprenyl diphosphate synthase (DPS) of *M*.

*tuberculosis* (32) and zFPPS(31). In the structure of DPS, the extreme C terminus of one monomer of decaprenyl diphosphate synthase interacts with the active site of the other subunit. Based on our biochemical data and the available structural information for homomeric *cis*PTs, it is likely that Arg^290^ in the RxG motif of NgBR is involved in IPP binding and the flexible nature of glycine in the motif, may permit conformational changes to seal the active site cavity. Collectively these data imply that both subunits contribute to enzymatic activity and the C-terminal tail of NgBR regulates aspects of *cis*PT catalytic activity.

The general function of NgBR orthologs as a *cis*PT subunit is further supported by the heteromeric *cis*PT from *M. acetovorans*. MaUPPS-B (NgBR ortholog) is missing any predicted TM domains and yet MaUPPS-A/ MaUPPS-B complex is indispensable for supporting the growth of triple deletion strain of *S. cerevisiae*. Since the mutations in conserved region of MaUPPS-B have similar impact on MaUPPS activity as those in distantly related h*cis*PT, we postulate a common mechanism for both enzymes.

In summary, this study strongly advances the concept that eukaryotic *cis*PT is composed of two subunits, NgBR and hCIT, and this two component system is conserved in *Methanosarcina acetovorans*. Interestingly, conservation of the RxG motif across phyla implicates a catalytic role of NgBR orthologs in the synthesis of polyprenol diphosphates critical for cellular function. Future structural studies on the heteromeric NgBR/hCIT complex will permit a deeper mechanistic understanding of how the C terminus regulates substrate binding and catalysis.

## Materials

Unless otherwise stated, all reagents were of analytical grade and purchased from Sigma Aldrich (St. Louis MO), Thermo Fisher Scientific (Waltham, MA), Zymo Research (Irvine, CA). Restriction enzymes were from New England Biolabs (Ipswich, MA). [1-^14^C]-IPP (50 mCi/mmol) was purchased from American Radiolabeled Chemicals (St. Luis, MO). Reverse phase thin layer chromatography (RP18-HTLC) plates were from Merck (Darmstad, Germany). Dolichol 19 standards was purchased from American Radiolabeled Chemicals (St. Luis, MO) and undecaprenol was obtained from Isoprenoids Collection of Institute of Biochemistry and Biophysics, PAS. Primary antibodies used in this study include *α-NgBR* (Abcam, ab168351), *α-* DHDDS (Sigma, SAB2100572), *α* -Strep-tag (IBA Solutions for Life Sciences, StrepMAB-Classic). Site-directed mutagenesis was performed using TagMaster Site-Directed Mutagenesis Kit (GM Biosciences). The *Escherichia coli* UPPS was amplified from genomic DNA. *Methanosarcina acetovorans* ORFs encoding MaUPPS-A (MA3723) and MaUPPS-B (MA4402) were amplified form synthetically synthetized codon optimized for expression in *S. cerevisiae* gBlocks^®^ Gene Fragments (ITD integrated Technologies, Coralville, Iowa). Gl*cis*PT, NgBR and hCIT isoform 1 were amplified from previous plasmids (REF). The cassette containing synthetic intron and internal ribosome entry site (IRES) was amplified form piRESneo1 (Clontech). Constructs used in this study are listed in Table I-S. cloning primers are listed in Table 11-S and mutagenesis primers are listed III-S. Invitrogen Gateway Cloning strategy was used to insert cDNA into yeast expression vectors. All PCR products were cloned into pCR8/GW/TOPO TA (Invitrogen), sequenced and sub-cloned into final yeast expression vector pKG-GW1 or pKG-GW2. To express His-Strep tagII-Gl*cis*PT in bacteria, the Strep tagII-Gl*cis*PT was amplified with restriction enzyme recognition sites and Strep -tag sequence inserted in the primers. The PCR product was ligated in frame with internal 6 His tag of pRSF-DUET1 plasmid.

To express both subunits of h*cis*PT from single mRNA in mammalian cells, the internal ribosome entry site (IRES) surrounded by two multicloning sites was introduced into pCEP4 (Invitrogen) using NEBUilder HiFi DNA Assembly (NEB) to obtain the pKGmDUET vector (details Supplemental Method). 6-His internally tagged NgBR was sub-cloned into BamH I/Not I sites of pKGmDUET from pKG-GW2-(31-HIS)NgBR plasmid and SmaI/EcoRI/ Klenow treated Strep-hCIT was sub-cloned into PmeI site from pKG-GW-Strep-hCIT plasmid.

### Yeast complementation assay

For yeast complementation analysis of *cis*PTs, *S. cerevisiae* strains KG405 *(nus1Δ rer2Δ srt1Δ)*, carrying the Gl*cis*PT gene on a plasmid with a *URA3* marker was used (43). To analyze homomeric Gl*cis*PT and EcUPPS mutants, strain KG405 was transformed with vector pKG-GW1 (leucine selection) carrying WT or mutated versions of corresponding genes, or empty vector as negative control. To phenotypically analyze h*cis*PT, strain KG405 was transformed with vectors pKG-GW1 carrying hCIT variants (leucine selection) and pKG-GW2 carrying NgBR variants (methionine selection) in combination, or empty vectors as negative control. To analyze putative heteromeric UPPS of *Methanosarcina acetovorans* (MaUPPS), strain KG405 was transformed with vector pKG-GW1 carrying MaUPPS-A (MA3723, hCIT ortholog) and vector pKG-GW2 carrying WT and mutated variants of MaUPPS-B (MA4402, NgBR ortholog) in combination or with the corresponding empty vectors as negative control.

Transformed yeast cells were grown overnight at 30°c in synthetic defined medium lacking uracil and leucine or uracil, methionine and leucine were streaked onto synthetic defined medium containing all amino acids, nucleotide supplements and 1% (w/v) 5-FOA (Zymo Research) and onto YPD plates. The plates were incubated for up to 7 days at 30°C. Colonies growing on the 5-FOA plates were streaked on synthetic defined medium lacking uracil and incubated at 30°c for 3 days to verify the loss of the pNEV-GlcisPT plasmid.

Yeast strain KG405 and its derivative carrying MaUPPS complex, h*cis*PT complex or single subunits enzymes expressed from pKG-GW1 plasmid instead of pNEV-GlcisPT were cultured in 2% (wt/vol) Bacto peptone and 1% (wt/vol) yeast extract supplemented with 2% glucose (wt/vol) (YPD). Synthetic minimal media were made of 0.67% (wt/vol) yeast nitrogen base and 2% (wt/vol) supplemented with auxotrophic requirements. For solid media, agar (BD, Sparks MD) was added at a 2% (wt/vol) final concentration. Yeast cells were transformed using the Frozen-EZ Yeast Transformation II Kit (Zymo Research).

### Purification of human *cis*PT

To purify the human hcisPT complex, constructs containing internally tagged 6His-NgBR and N-terminally tagged Strep-hCIT(pKGmDUET-hCIT/NgBR) were transiently transfected in 200 ml culture of Expi293F cells according to the manufacturer’s protocol (Invitrogen). Cells were harvested 72 hr post post-transfection by centrifugation, and washed with PBS. Each gram of the cells was re-suspended in S-tactin buffer (100 mM Tris HCL, pH 8, 300 mM NaCl, 2 mM 2-mercaptoethanol, 1 mM MgCl_2_) and cells were disrupted by sonication on ice. Unbroken material was cleared with a 1000 g centrifugation, and the supernatant from this spin was re-centrifuged at 200 000 g for 30 min at 4°C to obtain total membrane fractions. Total membrane fraction was homogenized in Stactin buffer supplemented with 0.5% Triton X100 to solubilize membrane proteins. Solubilization was followed by additional 200,000 x g centrifugation. Strep-hCIT was purified from 200,000 g supernatant using the *Strep*-Tactin XT system (IBA GmbH). Strep-hCIT/6-His-NgBR complex was eluted from *Strep*-Tactin XT resin with 100 mM Tris HCL, pH 8, 300 mM NaCl, 2 mM 2-mercaptoethanol, 1 mM Mgcl_2_, 0.1% Triton X 100, 5% glycerol, 50 mM biotin elution buffer. The Strep-Tactin purification of Strep-hCIT was followed by nickel column purification of 6His-NgBR using HisPur Ni-NTA resin (Thermo Fisher Scientific). Sephadex G-25 in PD-10 Desalting Columns (GE Healthcare Life Sciences) were used to exchange buffer for 20 mM NaPi, pH 8, 300 mM NaCl, 2 mM 2-mercaptoethanol, 1 mM Mgcl_2_, 0.1% Triton X 100, 20% glycerol. Purification efficiency was tracked by Western blot analysis of each fraction, Coomassie Staining of SDS PAGE of final eluate and measurement of specific *cis*PT activity (Figure 1 and Table 1).

### Purification of GlcisPT

To purify Gl*cis*PT, pRSF-DUET1- Gl*cis*PT plasmid was transformed into *E. coli* Rosetta 2 cells (Novagen). *E. coli* was grow in auto-induction medium (44) till the logarithmic growth phase at 37°C flowed 24 hr incubation at 17°C to express heterologous protein. Cells were harvested by centrifugation, washed with PBS and stored -80°C. The proteins were extracted using B-PER reagent (Thermo Fisher Scientific) supplemented with 1 mM Mgcl_2_ and 2 mM 2-mercaptoethanol. The Gl*cis*PT was first purified using nickel column HisPur Ni-NTA resin (Thermo Fisher Scientific) followed by the Strep-Tactin XF (IBA GmbH) purification. Sephadex G-25 in PD-10 Desalting Columns (GE Healthcare Life Sciences) were used to exchange buffer for 20 mM NaPi, pH 8, 300 mM NaCl, 2 mM 2-mercaptoethanol, 1 mM Mgcl_2_, 0.1% Triton X 100, 20% glycerol. Purification efficiency was tracked by Western blot analysis and Coomassie Staining of SDS PAGE of final eluate.

### Size Exclusion Chromatography

The size exclusion chromatography was carried out using an ÄKTA^TM^ purifier (GE Healthcare Life Sciences, Pittsburgh, PA, USA) with the size exclusion column Superdex^TM^ 200 10/300 GL (GE Healthcare Life Sciences) at flow rate of0.5 ml/min. The UV absorption at 254 nm was measured as a readout for protein elution. The used buffer contained 50 mM TrisHCl buffer, pH 8.0 and 300 mM NaCl, 2 mM 2-mercaptoethanol, 1 mM Mgcl_2_, 0.1% Triton X100, 5 % (v/v) glycerol. The column was calibrated by using the LMW- and HMW-calibration kit (GE Healthcare Life Sciences). For determination of the void volume Blue Dextran (2,000 kDA; GE Healthcare Life Sciences) was used. Size determination was calculated based on the standard linear equation based on the calibration of the column.

### cisPT enzymatic activity of hcisPT

Standard incubation mixture contained, in a final volume of 50 μl, 50 μM FPP,100 μM [1- 14C]-IPP (55 mCi/mmol) 50 mM Tris-HCl pH 8, 1 mM Mgcl_2_, 20 mM 2–mercaptoethanol, 1mg/ml BSA, 10mM KF and 1% (w/v) phosphatidylinositol. Membrane or crude protein (100 μg) or 20-100 ng of purified protein was used for activity assays. In some experiments with crude membranes, zaragozic acid A (10 μM) was added. Reactions were incubated for 60 min at 37°C and terminated by the addition of 1 ml of chloroform-methanol 3:2. The protein pellet was removed by centrifugation and the supernatant was washed three times with 115 volume of 10 mM EDTA in 0.9% NaCl. The incorporation of ^14^C-IPP into organic fractions containing polyprenyl diphosphate was measured by scintillation counting. To analyze the length of polyprenol diphosphates lipids were subjected to mild acid hydrolysis (1 hr incubation at 80 °C in 500 μl of N HCL solution). Dephosphorylated lipids were extracted 3 times with 1 volume of hexane. Organic fraction was washed with **1** volume of water. Then organic solvent was evaporated and lipids were loaded together with dolichol19 as internal standard onto HPTLC RP-18 plates with and run in acetone containing 50 mM H_3_P0_4_. Plates were exposed to film to visualize the products of IPP incorporation.

### Kinetics parameters

Standard 25 μl or 100 μl (hCIT/ NgBR^G292^ mutant) reaction mixture containing 9–25nM h*cis*PT was used. To measure kinetic parameters for FPP, 0.1-50 μM FPP was use along with 100 μM IPP for wild type enzyme, hCIT^K42E^ and NgBR^H100A^mutants, 200 μM IPP for NgBR^R292H^ mutant and 400 μM IPP for NgBR^G292^ mutant. To measure kinetic parameters for IPP, 0.1-400 μM IPP was use along with 50 μM FPP. The initial velocity data were fitted to Michaelis-Menten equation using the GraphPad Prism 7.02 computer program (GraphPad Software, Inc.) to obtain *Km* values. k_cat_ values were obtained from Michaelis-Menten equation for IPP.

### cisPT enzymatic activity of GlcisPT

Gl*cis*PT activity was measured as before((41) with minor modifications. Briefly the incubation mixture contained, in a final volume of 50 μ,l, 45 μM FPP,100 μM [1- 14C]- IPP (55 mCi/mmol) 25 mM Tris-HCl pH 7.4, **1** mM Mgcl_2_, 20 mM 2–mercaptoethanol, 10 mM KF, 0.1% Triton X100, 1 mg/ml BSA and 2 μg of purified enzyme. After 60 min incubation at 37°, the reaction was terminated by the addition of 1 ml of chloroform-methanol 3:2. The protein pellet was removed by centrifugation and the supernatant was washed three times with 115 volume of 10 mM EDTA in 0.9% NaCl. The incorporation of 14C-IPP into organic fractions containing polyprenyl diphosphate was measured by scintillation counting. To analyze the length of polyprenol diphosphates the 200 μl, reaction was stopped by addition of HCL to obtain 1N final concentration. Lipids were subjected to mild acid hydrolysis, extracted with hexane and loaded onto HPTLC RP-18 together with undecaprenol as internal standard. After running in acetone containing 50 mM H_3_P0_4_ plates were exposed to film to visualize the products of IPP incorporation.

### cisPT enzymatic activity of MaUPPS

For *S*. *cerevisiae* expressing MaUPPS complex membrane fractions were prepared as described (45)and *cis*PT activity measured (45,46) with minor modifications. Briefly the incubation mixture contained, in a final volume of 100 μl, 45 μM FPP,100 μM [1- 14C]- IPP (55 mCi/mmol) 25 mM Tris-HCl pH 7.4, 1 mM MgCh, 20 mM 2–mercaptoethanol, 10 μM KF, 10 μM Zaragozic acid A and 250 μg of membranes protein. After 90 min incubation at 30°C, the reaction was terminated by the addition of 4 ml of chloroform-methanol 3:2. The protein pellet was removed by centrifugation and the supernatant was washed three times with 115 volume of 10 mM EDTA in 0.9% NaCl. The organic phase was concentrated under a stream of nitrogen. Then organic solvent was evaporated and lipids were loaded onto HPTLC RP-18 precoated plates with a concentrating zone and run in acetone containing 50 mM H_3_P0_4_. Plates were exposed to film to visualize the products of IPP incorporation. To measure incorporation of radioactive IPP into polyprenol fraction, the gel from the zone containing radiolabeled polyprenols was scraped and subjected to liquid scintillation counting. To analyze the length of polyprenol diphosphates before running the TLC analysis lipids were subjected to mild acid hydrolysis. Lipids were loaded onto HPTLC RP-18 together with undecaprenol as internal standard and subjected to radio-autography as was described above.

## Acknowledgments

This work was supported by Grants ROI HL64793, ROI HL6137l and HLI33018 from the National Institutes of Health and the Leducq Fondation (MIRVAD network) to WCS and an American Heart Association Scientist Development Grant to EJP. We acknowledge the gift of undecaprenol from Dr. Ewa Kula-Swiezewska (Polish Academy of Science).

---

**Figure 4 – figure supplement 1.**
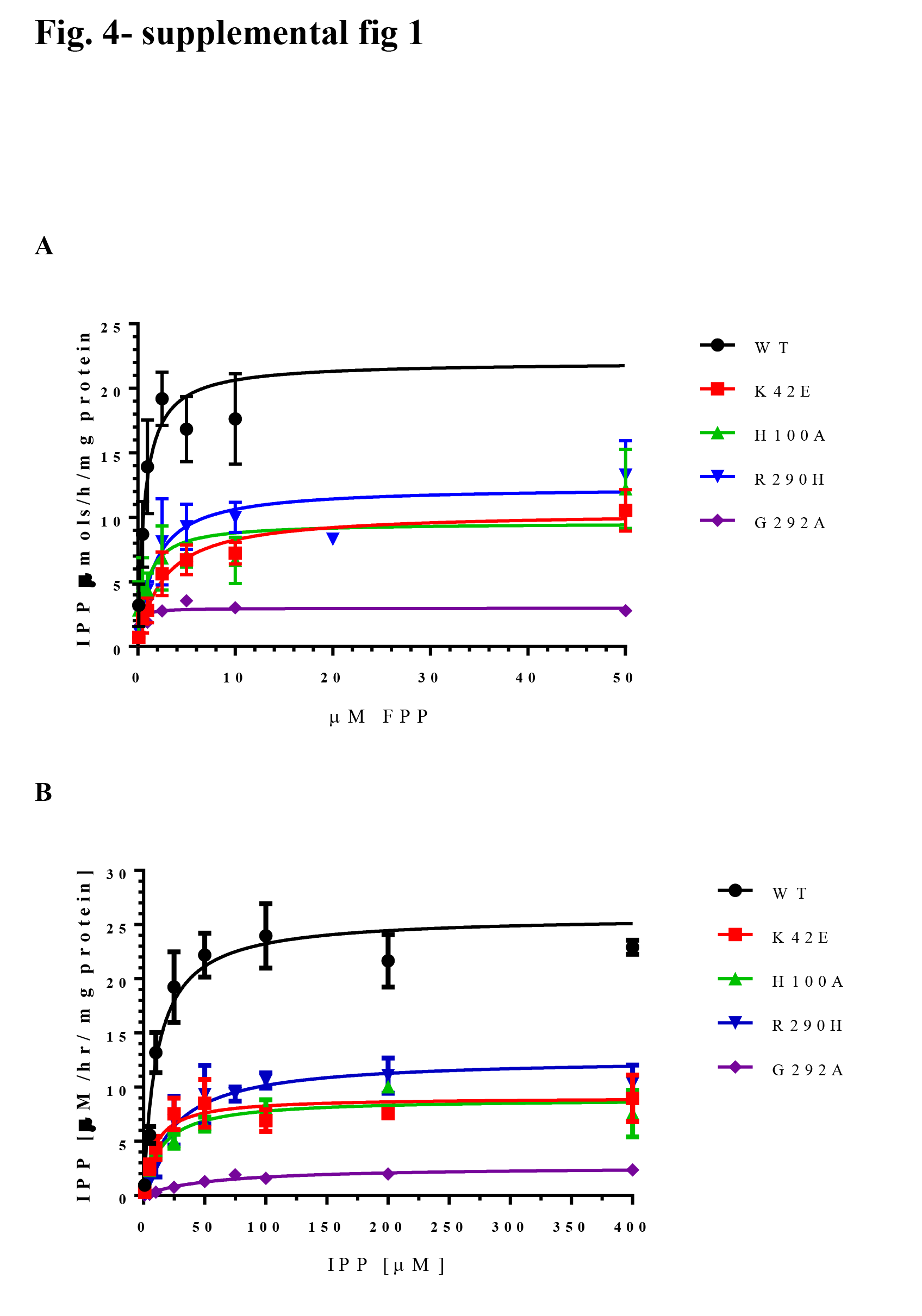
Michaelis-Menten plot of hcisPT kinetics. **A** The measurements of Km for FPP and B. IPP. Each data points represent 3–8 independent measurements (n=1 for measurement of Km for FPP of hCIT/NgBR^G292A^ enzyme, and n=2 for other samples)‥ Details are provided under “Materials and Methods.”.

